# Single-cell analysis of chromatin silencing programs in development and tumor progression

**DOI:** 10.1101/2020.09.04.282418

**Authors:** Steven J. Wu, Scott N. Furlan, Anca B. Mihalas, Hatice S. Kaya-Okur, Abdullah H. Feroze, Samuel N. Emerson, Ye Zheng, Kalee Carson, Patrick J. Cimino, C. Dirk Keene, Jay F. Sarthy, Raphael Gottardo, Kami Ahmad, Steven Henikoff, Anoop P. Patel

## Abstract

Single-cell analysis has become a powerful approach for the molecular characterization of complex tissues. Methods for quantifying gene expression^1^ and chromatin accessibility^2^ of single cells are now well-established, but analysis of chromatin regions with specific histone modifications has been technically challenging. Here, we adapt the recently published CUT&Tag method^3^ to scalable single-cell platforms to profile chromatin landscapes in single cells (scCUT&Tag) from complex tissues. We focus on profiling Polycomb Group (PcG) silenced regions marked by H3K27 trimethylation (H3K27me3) in single cells as an orthogonal approach to chromatin accessibility for identifying cell states. We show that scCUT&Tag profiling of H3K27me3 distinguishes cell types in human blood and allows the generation of cell-type-specific PcG landscapes from heterogeneous tissues. Furthermore, we use scCUT&Tag to profile H3K27me3 in a brain tumor patient before and after treatment, identifying cell types in the tumor microenvironment and heterogeneity in PcG activity in the primary sample and after treatment.

---

Significant portions of the genome are actively repressed to create barriers between cell type lineages during development^4^. In particular, trimethylation on lysine 27 of histone H3 (H3K27me3) in nucleosomes by PcG proteins is crucial for gene silencing during normal differentiation and thus for maintaining cell identity^5^. Conversely, derangements in PcG silencing permit aberrant gene expression and disease^6^. Therefore, methods for assaying silenced chromatin can provide insights into a variety of processes ranging from normal development to tumorigenesis.

Scalable methods for assessing silenced chromatin at the single-cell level have not been widely available. We set out to use chromatin profiling of single cells to assess gene silencing and to develop a framework for analysis. Our approach builds on Cleavage Under Targets and Tagmentation (CUT&Tag), which uses specific antibodies to tether a Tn5 transposome at the sites of chromatin proteins in isolated cells or nuclei. Activation of the transposome then tagments genomic loci with adapter sequences that are used for library construction and deep sequencing, thereby identifying binding sites for any protein where a specific antibody is available^3^. Our earlier work demonstrated that CUT&Tag profiling of the H3K4me2 histone modification efficiently detected gene activity, much like ATAC-seq, while H3K27me3 profiling detected silenced chromatin that may be epigenetically inherited^3^.

To determine whether single-cell chromatin landscapes were sufficient to distinguish different cell types, we performed CUT&Tag on H1 human embryonic stem cells (H1 hESCs) using an anti-H3K27me3-specific antibody in bulk and then distributed single cells for PCR and library enrichment on the ICELL8 system (Fig. 1a). We compared this to previously published H3K27me3 scCUT&Tag profiles of K562 cells and hESC^3^ to determine whether standard approaches to single-cell clustering could distinguish cell types based on H3K27me3 signal. As PcG domains typically span >10 kilobases, we grouped read counts in 5 kilobase bins across the genome and used this for latent sematic indexing (LSI) based dimensional reduction and UMAP embedding, followed by standard Louvain clustering using the ArchR package^7^ (see methods). After quality control filtering (see methods, Supplementary Fig. 1a-g), UMAP embedding clearly separated 100% of 804 hESC cells with a median of 375 unique fragments from 908 K562 cells with a median of 6064 unique fragments independent of batch effects (Fig. 1b). Interestingly, hESC had 6% of the number of unique fragments when compared to K562 cells (Supplementary Fig. 1f). This demonstrates that stem cells have lower global H3K27me3 levels than more differentiated cell types^8^. Despite down-sampling the number of unique fragments per cell to the same median value for both datasets, H3K27me3 signal still readily distinguished the two cell types (Supplementary Fig. 1g) confirming that clustering was driven by differences in H3K27me3 signal and not number of unique fragments.

**Figure 1:**
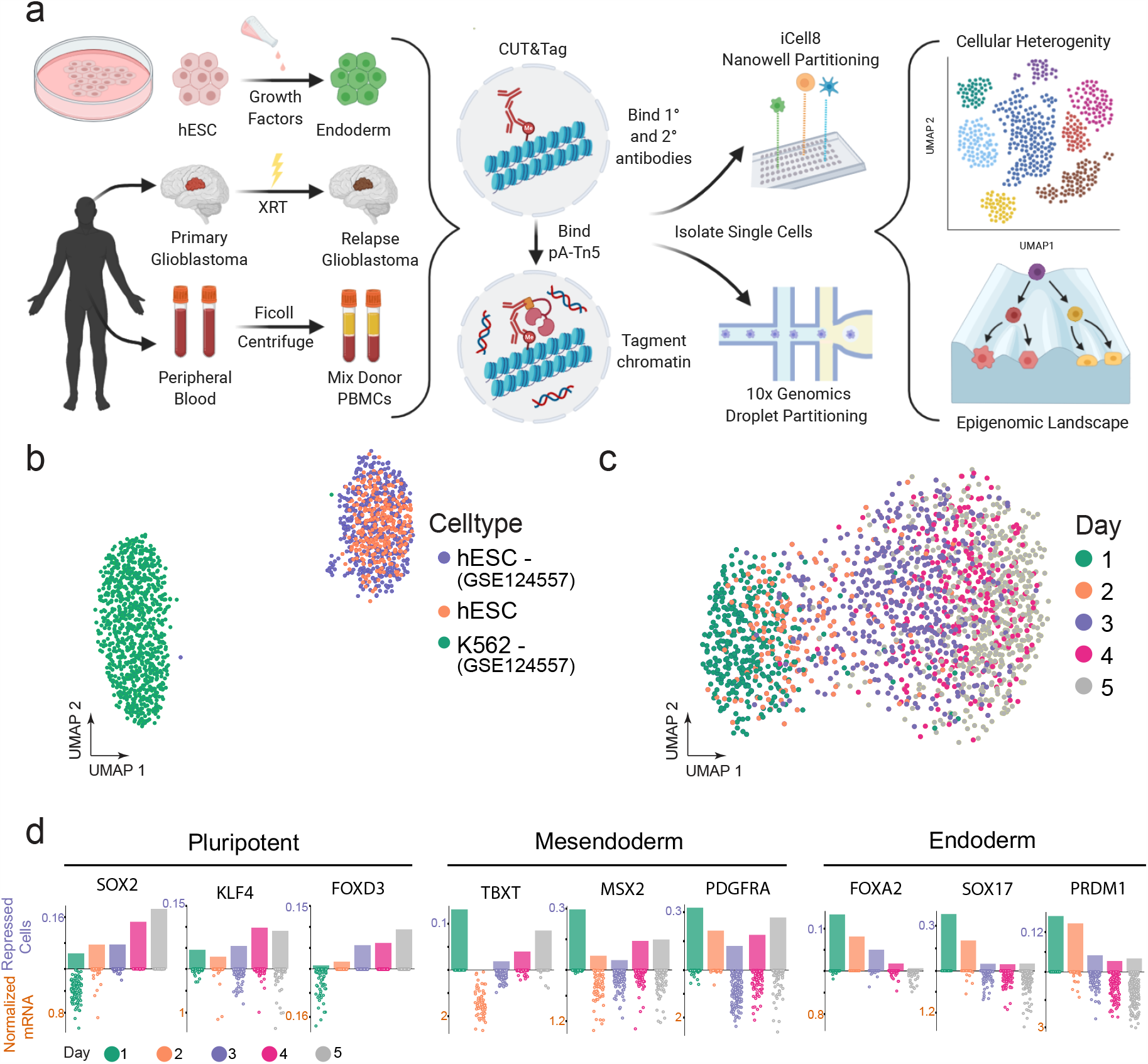
Single-Cell CUT&Tag resolves distinct cell-types and maps repressive chromatin domains in early hESC development. a) Schematic of scCUT&Tag applied to nuclei isolated from cell culture, a model endoderm differentiaton system, blood cells, and a human brain tumor. Single cells are then partitioned using either the 10X Genomic or iCELL8 microfluidic systems. b) UMAP embedding of scCUT&Tag for a repressive histone modification, H3K27me3, in K562 (n=908) and hESC (n=804) single cells. c) UMAP embedding of scCUT&Tag for a repressive histone modification, H3K27me3, in a 5 day differentiation time course from hESC to definitive endoderm (total n=1830). Cell types are colored according to the day along the time course in which they were harvested. d) A bar plot representing the percent of single cells that are repressed at each specific gene. The jitters below depict scRNA-seq for the same timepoint. The upper axis corresponds to scCUT&Tag (percent of single cells repressed) and lower axis corresponds to scRNA-seq (normalized mRNA counts). From left to right, well-known TF markers for pluripotent, mesendoderm, and definitive endoderm cells.

Cellular determination and differentiation proceed by a controlled sequence of gene activation and gene repression. To study gene silencing during development, we differentiated hESCs towards definitive endoderm^9^. We confirmed differentiation by immunofluorescence staining of stage-specific transcription factors (Supplementary Fig. 2a). UMAP embedding of 1830 scCUT&Tag H3K27me3 profiles with a median of 279 fragments revealed a developmental trajectory, independent of batch effect (Supplementary Fig. 2b), from hESC to definitive endoderm (Fig. 1c) that was punctuated by stem-like states on days 1-2 followed by a rapid progression towards differentiation on days 3-5. To determine if changes in chromatin silencing corresponded to changes in gene expression, we examined known markers of stem cells and endoderm differentiation in single-cell aggregate profiles from each day. Overall, H3K27me3 signal at a marker gene was inversely correlated with expression based on a published scRNA-seq dataset^9^. Stem cell markers such as SOX2, KLF4, and FOXD3 are expressed in hESCs and lack H3K27me3 but are silenced as differentiation proceeds (Fig. 1d). Between day 2 and 3, hESCs transition into a mesendoderm state (characterized by expression of TBXT, MSX2, and PDGFRA) in which they have the developmental potential to either become mesoderm or endoderm^9^. This is illustrated in our data between day 2-3 where chromatin silencing at mesoderm markers is lower (Fig. 1d). As differentiation proceeds, endoderm markers such as FOXA2, SOX17, and PRDM1 become active and lose H3K27me3 signal (Fig. 1d). Finally, markers of ectoderm (PAX6 and LHX2), are not expressed, and accumulated H3K27me3, consistent with silencing of these loci (Supplementary Fig. 2c). Pseudo-temporal ordering of single cells recapitulated our real-time results (Supplementary Fig. 2d).

Having established that scCUT&Tag readily identifies dynamic changes in chromatin silencing, we next sought to determine whether chromatin profiles could distinguish cell types in a more complex tissue. To do so, we adapted scCUT&Tag to the 10X Genomics microfluidics platform and profiled H3K27me3 in mixed peripheral blood mononuclear cells (PBMCs) collected from two healthy donors. Briefly, we performed scCUT&Tag in bulk on 1 million cells and then loaded two lanes of a 10X Genomics microfluidic chip with 10,000 nuclei each to obtain technical replicates (Supplementary Fig. 3). We implemented a ‘chromatin silencing score’ (CSS), which uses the gene activity score (GAS) model in ArchR^7^ to create a proxy for the overall signal associated with a given locus. Quality control filtering resulted in 9,917 cells with a median of 1,110 unique fragments per cell for which we performed dimensionality reduction and embedding as described above (Fig. 2a). The median number of reads falls in the range expected for cell type variation, in spite of the platform differences in our study.

**Figure 2:**
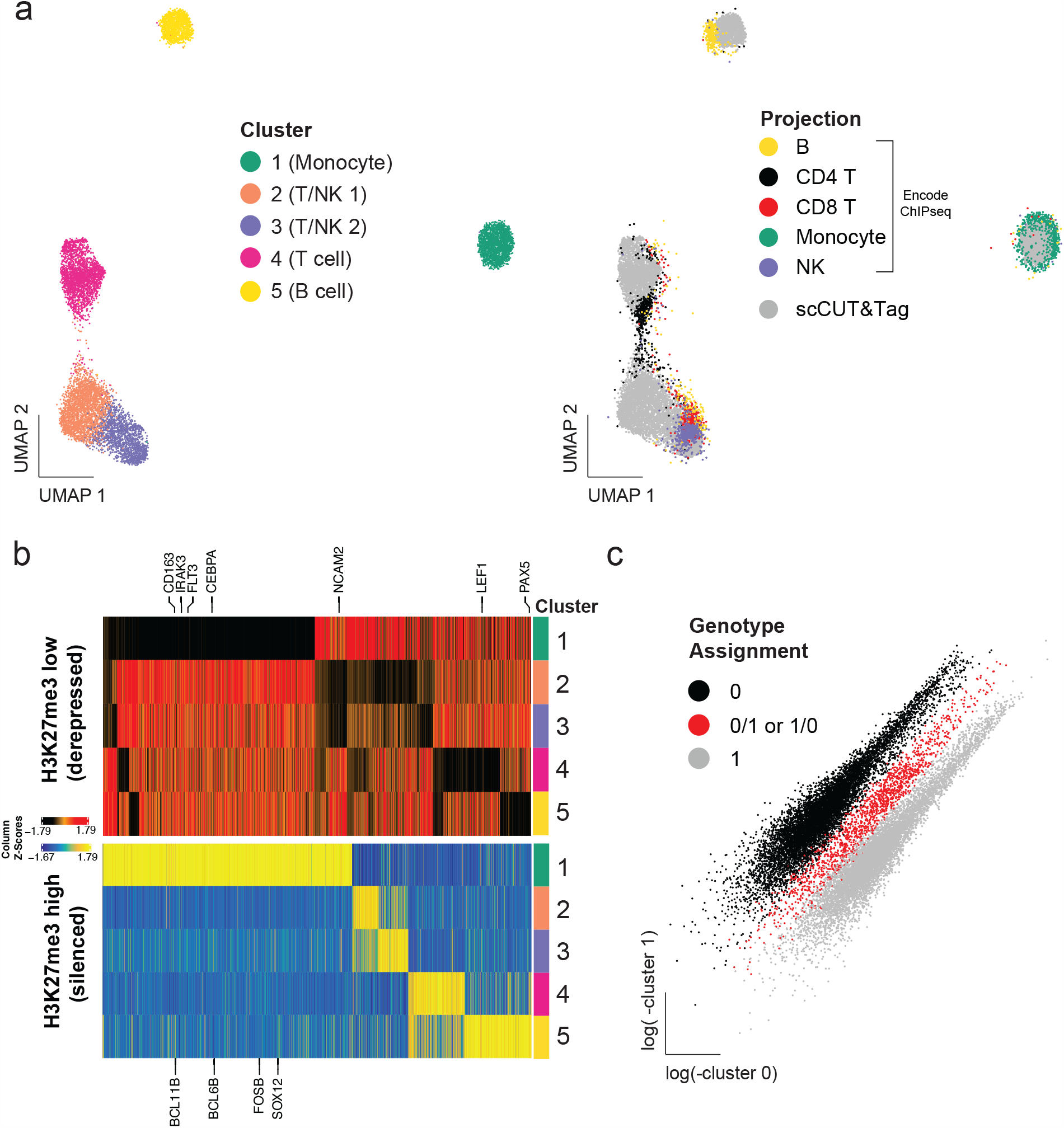
scCUT&Tag for H3K27me3 readily identify major subtypes in PBMC. a) Left - UMAP embedding of single cell data from PBMC. Unsupervised clustering revealed 5 clusters. Right - UMAP projection of downsampled ChIPseq bulk data from primary sorted bulk datasets for major PBMC cell-types (see Supplementary methods for GSE citations) on single cell CUT&Tag data on left. b) Heatmap of genes with significantly low (top) or high (bottom) H3K27me3 signal in each cluster (row). Fold change < −2 (top) or > 2 (bottom); q-value < 0.05 (both). Cell type specific genes are highlighted. c) Sparse mixture model clustering (using souporcell) of genotype variant calls from the PBMC data colored by genotype assignment (prior to multiplet removal).

We then set out to identify the major cell types in the data using two methods. We first down-sampled publicly available bulk H3K27me3 ChIP-seq data (ENCODE) and used the UMAP transform function to “project” the ChIP-seq data onto our UMAP embedding as previously described^10^ (Fig. 2a). We used the CSS score to identify cell-type specific marker genes that showed a lack of H3K27me3 enrichment because active genes will have a low CSS. Therefore, we would expect a low CSS for a cell type specific marker gene in the cluster that corresponded to that cell type (Fig. 2b). Overall, cluster identification by CSS annotation matched our assignments by ChIP-seq projection (Fig. 2a) and distinguished major cell types in unsorted PBMCs including those of lymphoid (T cell, NK cell, B cell) and myeloid lineages (monocyte). We recovered the proportions of major cell types within the range of normal adult blood (Supplementary Table 1). Using this method, we can therefore generate cell type specific PcG landscapes across heterogenous cell types within a sample, obviating the need for physical cell sorting and minimizing confounding effects of batch effect, read depth, or sample heterogeneity (Supplementary Fig. 3). This allowed us to identify the top differentially PcG-silenced loci across the major cell types in PBMCs (Fig. 2b). We also profiled PBMCs with the active mark H3K27ac and recovered the major cell types in a similar proportion as H3K27me3 scCUT&Tag (Supplementary Fig. 4, Supplementary Table 1).

We next demultiplexed each biological donor using Souporcell. In brief, the algorithm identifies genotypic differences between single cells by variant calling aligned reads^11^. The variant calls can also be used to identify multiplets. Using this method we were able to differentiate cells from each donor (Fig. 2c). Clustering was not driven by donor specific effects but rather by cell type differences (Supplementary Fig. 3b).

Having established scCUT&Tag can profile developmental systems and heterogenous tissues, we used scCUT&Tag to interrogate PcG based clustering in glioblastoma (GBM), a human central nervous system tumor that is known to have a heterogeneous microenvironment^12^, exhibit intratumoral heterogeneity^13^ and have pseudo-hierarchical organization that mimics development^12, 14, 15^. In this tumor type, changes in PcG chromatin silencing can mediate emergence of resistant cell populations^16^.

We profiled H3K27me3 in 1,311 single nuclei (3,643 median fragments/cell) using the 10X scCUT&Tag workflow from a primary glioblastoma which had been snap-frozen shortly after surgical removal. We distinguished four major cell populations within the sample (Fig. 3a). To annotate clusters, we constructed CSS of previously-defined marker loci^12^, and annotated clusters that correspond to microglia (Cluster 1, low CSS at the PTPRC gene), neurons (Cluster 3, low CSS at RBFOX3), oligodendrocytes (Cluster 4, low CSS at MOBP), and other neural lineage cells, including tumor cells (Cluster 4, low CSS at SOX2) (Fig. 3b). To confirm cluster annotations, we projected CUT&RUN bulk data from a glioma stem cell line (UW7gsc) derived from the same patient, two established neural stem cell lines (U5 and CB660)^17^, and ENCODE^18^ ChIP-seq bulk data for monocytes (proxy for microglia) and astrocytes. Projection onto the scCUT&Tag tumor sample embedding confirmed CSS annotations (Fig. 3c). UW7gsc projected to the center of the largest cluster, presumably made up of tumor cells. The astrocyte data projected to a smaller satellite cluster within the neural lineage cells. The neural stem cell line data localized to both the tumor cell cluster as well as the astrocyte cluster (Supplementary Fig. 5). This may reflect spontaneous differentiation of neural stem cells towards the astrocyte lineage *in vitro* or reflect subtle changes in cell state such as lineage priming^19^.

**Figure 3:**
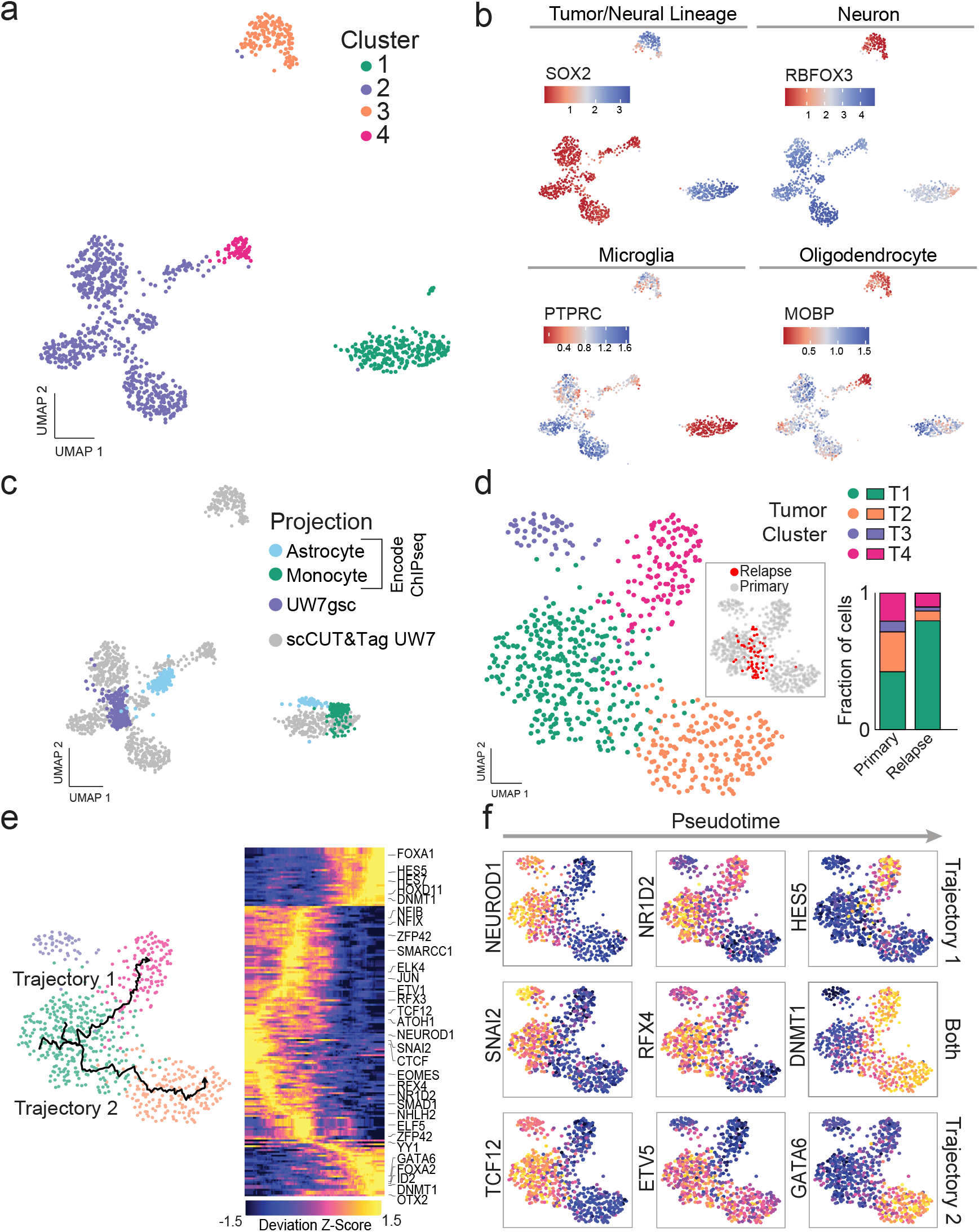
scCUT&Tag data for H3K27me3 for a human glioblastoma primary and relapse sample demonstrates heterogeneity in PcG distribution within tumor cell clusters and cluster enrichment after treatment. a) UMAP embedding of single cells from a primary human glioblastoma based on H3K27me3 signal. b) Cluster annotation using chromatin silencing scores for key markers genes identifies microglia (PTPRC), neurons (RBFOX3), oligodendrocytes (MOBP), and tumor cells (SOX2). c) UMAP transform and projection of bulk ChIP seq (monocytes, astrocytes) or bulk Cut&Run (UW7gsc) onto patient sample. d) Left - UMAP co-embedding of tumor cells from primary and relapse sample. Inset highlights locations of cells from relapse sample. Right - Barplot demonstrating fraction of cells in each sample (Primary, Relapse) that belong to each cluster. e) Left - Two pseudotime trajectories starting with cluster T1 (presumed stem-like cluster) and ending in either cluster T4 (Trajectory 1) or cluster T2 (Trajectory 2). Right - Heatmap of 132 significant motif deviations based on H3K27me3 activity within peaks from aggregated tumor cell ATAC-seq data. Motif deviations are ordered by pseudotime. f) UMAP plots for tumor cells colored by deviation scores for selected motifs. Left column shows early motifs in pseudotime that are commonly silenced including NEUROD1, SNAI2, and TCF12. Middle column shows silenced programs that diverge according to trajectory (NR1D2 in Trajectory 1 and ETV5 in Trajectory 2) or are common across trajectories (RFX4). Right column shows silenced programs specific to terminal pseudotime for Trajectory 1 (HES5), Trajectory 2 (GATA6) or both (DNMT1).

To understand how the tumor changed with treatment, we performed scCUT&Tag profiling for H3K27me3 for a relapse sample obtained via rapid autopsy from the patient 5 months after surgery and radiation therapy. Application of quality control metrics followed by low dimensional embedding identified 4 distinct cell types in the relapse sample (Supplementary Fig. 6a). Projection of the 1168 autopsy single cell profiles (16,232 median fragments/cell) onto the primary tumor UMAP embedding allowed cell type identification, including 71 cells that colocalized to the tumor cell cluster (Supplementary Fig. 6b).

We focused on the tumor cells, co-embedding the 71 relapse tumor single cells with the 640 primary tumor single cells. After batch correction, we identified 4 clusters within the tumor cell data with distinct H3K27me3 profiles (Fig. 3d - left). Examining the distribution of cell states across the two timepoints, we noted an enrichment for Cluster T1 in the relapse specimen (Fig. 3d - right). The relapse tumor cells had higher background signal when compared to the primary tumor cells as determined by FRiP analysis (Supplementary Fig. 6c). To further confirm that the relapse cells were most similar to Cluster T1, we characterized reads in relapse cells that were present in genomic regions that most significantly distinguished the primary tumor clusters (Supplementary Fig. 6d). This analysis confirmed similarity of the relapse tumor cells to Cluster T1. Gene set enrichment analysis using the CSS matrix identified potential programs silenced (positive enrichment scores) and derepressed (negative enrichment score) in this cluster. Interestingly, the Verhaak_glioblastoma_proneural gene set appears to be silenced in the resistant cell cluster (Supplementary Fig. 7), consistent with the idea that tumor evolution may induce a proneural-to-mesenchymal shift^20^. In contrast, low CSS was observed at gene sets with high CpG content that are marked by H3K27me3 in whole brain^21^. The lack of H3K27me3 signal in this tumor cluster suggests that the PcG landscape of glioblastoma cells resembles a stem-like state rather than a terminally differentiated state^22^.

We next wanted to understand the relationship between the cell clusters. Clusters T1, T2 and T4 exist along a continuum, whereas cluster T3 is separated from the main tumor cell group. We focused on whether transcription factor programs are differentially silenced across clusters T1, T2, and T4. H3K27me3 domains are broad, spanning 10-100kb and covering many genes, enhancers, promoters, and intervening regions. Therefore, to limit motif searching to potential regulatory elements within H3K27me3 domains, we used single-cell ATAC-seq data (Supplementary. Fig 8) to annotate enhancers and promoters in tumor cell sub-clusters based on accessible chromatin. We then calculated TF motif enrichments and depletions in this set of curated genomic regions based on H3K27me3 signal. We examined motif deviations ordered over two pseudotime trajectories that started with cluster T1 (presumed stem-like cluster) and ended in either cluster T4 (Trajectory 1) or cluster T2 (Trajectory 2) (Fig. 3e - left). Motif deviations (n=132) were ordered according to pseudotime, identifying silenced motifs that spanned cluster T1 to cluster T2 and cluster T4 (Fig. 3e - right). At the apex of the trajectories, motif silencing was shared and included motifs for TFs such as NEUROD1, SNAI2, and TCF12 (Fig. 3f, left column). At intermediate pseudotime points, there were silenced motifs specific to Trajectory 1 (NR1DA2) or Trajectory 2 (ETV5) or shared by both (RFX4) (Fig. 3f, middle column). As pseudotime proceeds, Trajectory 1 showed evidence of HES5 motif silencing, while Trajectory 2 showed GATA6 motif silencing. Interestingly, the DNMT1 motif was strongly silenced across both pseudotime endpoints, concordant with the idea that PcG silencing of DNMT1 enriched promoters and enhancers is a common feature of differentiation^23^ (Fig. 3f, right column).

Fundamentally, we have shown here that repressive chromatin can be used to identify cell states *a priori* from heterogeneous normal and diseased tissues. This approach has far reaching applications, including generation of cell type specific chromatin atlases from archival tissue in a manner that does not require sorting of pure populations. We focused here primarily on a single chromatin mark, but this method can in theory be applied to any histone modification or DNA binding protein for which an antibody is available. As such, developing complete chromatin landscapes of complex tissues and disease states using scCUT&Tag will help decode the complex epigenetic machinery underlying gene expression. Broadly, our method for performing histone mark specific single-cell analysis adds to the growing list of single-cell ‘omic’ methods that can be used to understand heterogeneous cell populations.

## Materials & Methods

### Biological Material

H1 ES cells were purchased from WiCell (Cat#WA01-lot#WB35186). We used the following antibodies: Guinea Pig anti-Rabbit IgG (Heavy & Light Chain) antibody (Antibodies-Online ABIN101961), H3K27me3 (Cell Signaling Technology, cat #9733), H3K27ac (Millipore Sigma, cat# MABE647), SOX17 (R&D Systems, AF1924, Lot KGA0916121), OCT4 (Abcam, ab109183, lot gr120970-6), and H3K4me2 (Upstate 07–030, Lot 26335) The fusion enzyme, pA-Tn5 was generated as previously described^3^.

### hESC culture conditions

H1 ES cells were maintained on Corning Matrigel hESC-qualified Matrix, Corning (#354277), at 37C in mTeSR™1 from STEMCELL Technologies (Catalog #85850) with daily media replacement. When cell aggregates were 80% confluent, they were released using ReLeSR, STEMCELL Technologies (# 05872), per manufacturer’s instructions and incubated at 37°C for 3-5 minutes. Cells were released into a small volume of complete media by tapping of growth plate and aggregates reduced in size by gentle pipetting and passaged to desired ratio.

### hESC differentiation protocol

hESC were differentiated to definitive endoderm using the STEMdiff Definitive Endoderm Kit (cat #05110). The full protocol is available from STEMCELL Tech (https://cdn.stemcell.com/media/files/pis/29550-PIS_2_1_0.pdf?_ga=2.73376023.564267965.1597964514-138601152.1597964514). Briefly, hESC at 80% confluent were harvested using Gentle Cell Dissociation Reagent (STEMCELL Tech, cat #07174) and reseeded in a single-cell manner on Matrigel-plates. This was done daily for five days. Every 24 hours after a new differentiation culture was started and cells were incubated with DE differentiation medium according to the manufacture’s guideline. On the 5^th^ day, all five timepoints were harvested simultaneously using Accutase (STEMCELL Tech, cat# AT104-500). Immunofluorescence was used to confirm differentiation as previously described^24^.

### PBMC acquisition and processing

Healthy adult donors at the University of Washington underwent venipuncture and blood was collected using heparin-containing vacutainer tubes after consenting to participate in our study, Institutional Review Board protocol (#STUDY00008678). Additional PBMC specimens were obtained from consented donors at the Fred Hutchinson Cancer Research Center (IRB# 0999.209). Mononuclear cells were harvested from peripheral blood using gradient centrifugation. Cells were then washed twice with PBS and captured as outlined below.

### Brain tumor specimen acquisition, processing and culture

Adult patients at the University of Washington provided preoperative informed consent to take part in the study in all cases following approved Institutional Review Board protocols (IRB protocol #STUDY00002162). Fresh tumors were collected directly from the operating room at the time of surgery and either taken fresh or snap frozen immediately after removal in liquid nitrogen. Histopathologic diagnosis was confirmed by a board certified neuropathologist. Fresh tissue was enzymatically dissociated using a papain-based brain tumor dissociation kit (Miltenyi Biotec) as per manufacturer’s protocol. Cells were then cultured on laminin coated plates in DMEM/F12 supplemented with 1X N2/B27, 1% Penicillin/Streptomycin. Cultures were passaged as needed when confluent and considered stable after 3 serial passages. Cell line UW7gsc was used for this study at passage number 3. Autopsy tissue was collected with a post-mortem interval of approximately 8.75 hours after informed consent with a waiver from the University of Washington IRB. Tissue was snap frozen in liquid-nitrogen cooled isopentane. Tumor regions were sampled based on gross examination of brain sections and processed as outlined below.

### Nuclei preparation from brain tumor specimens

Frozen tissue was processed to nuclei using the ‘Frankenstein’ protocol from Protocols.io. Briefly, snap frozen tissue glioblastoma tissue was thawed on ice and minced sharply into <1 mm pieces. 500 ul chilled Nuclei EZ Lysis Buffer (Millipore Sigma NUC-101 #N3408) was added and tissue was homogenized 10-20 times in a Dounce homogenizer. The homogenate was transferred to a 1.5 ml Eppendorf tube and 1 mL chilled Nuclei EZ Lysis Buffer was added. The homogenate was mixed gently with a wide bore pipette and incubated for 5 minutes on ice. The homogenate was then filtered through a 70 um mesh strainer and centrifuged at 500g for 5 minutes at 4°C. Supernatant was removed and nuclei were resuspended in 1.5 mL Nuclei EZ lysis buffer and incubated for 5 minutes on ice. Nuclei were centrifuged at 500g for 5 min at 4°C. After carefully removing the supernatant (pellet may be loose), nuclei were washed in Wash Buffer (1x PBS,1.0% BSA,0.2 U/μl RNase Inhibitor). Nuclei were then centrifuged and resuspended in 1.4 ml Wash Buffer for two additional washes. Nuclei were then filtered through a 40 um mesh strainer. Intact nuclei were counted after counterstaining with Trypan blue in a standard cell counter.

### Chromatin Profiling: scCUT&Tag using the ICELL8 system/protocol

scCUT&Tag for the ICELL8 was carried out as previously described^3^. In brief, approximately 250,000 hESC (for each timepoint) were processed by centrifugation between buffer exchanges at 600xg for 3 minutes and in low-retention tubes. Cells were collected and washed with 1mL wash buffer (20 mM HEPES, pH 7.5; 150 mM NaCl; 0.5 mM Spermidine, 1× Protease inhibitor cocktail) at room temperature. Cells were incubated antibody diluted 1:50 in NP40-Digitonin Wash Buffer (0.01% NP40, 0.01% Digitonin in wash buffer) overnight. This wash buffer permeabilized the cells and released nuclei. Permeabilized nuclei were then rinsed once with NP40-Digitonin Wash buffer and incubated with anti-Rabbit IgG antibody (1:50 dilution) in 1 mL of NP40-Digitonin Wash buffer on a rotator at room temperature for 30 min. Nuclei were washed twice with NP40-Digitonin Wash buffer and incubated with 1:100 dilution of pA-Tn5 in NP40-Dig-med-buffer (0.01% NP40, 0.01% Digitonin, 20 mM HEPES, pH 7.5, 300 mM NaCl, 0.5 mM Spermidine, 1× Protease inhibitor cocktail) for one hour at RT on a rotator. Cells were washed 2x with NP40-Dig-med-buffer and resuspended in 150 µL Tagmentation buffer (10 mM MgCl2 in NP40-Dig-med-buffer) and incubated at 37 °C for 1 h. Tagmentation was stopped by adding 50 µL of 4× Stop Buffer (40.4 mM EDTA and 2 mg/mL DAPI) and the sample was held on ice for 30 min. Samples were then strained through a 10-micron cell strainer to remove clumps of cells.

The SMARTer ICELL8 single-cell system (Takara Bio USA, Cat. #640000) was used to array single cells previously described^3^. Briefly, cells were loaded onto a source plate and dispensed in to a SMARTer ICELL8 350 v chip (Takara Bio USA, Cat. # 640019) at 35 nanoliter per well. The chip was then spun down at 300xg for 5 minutes. Imaging on a DAPI-channel confirmed the presence of single-cells in specific wells. Non-single cell wells were excluded from downstream reagent dispenses. To index the whole chip, 72×72 i5/i7 unique indices (5184 micro-wells total) were dispensed at 35nL in wells that contained single cells followed by two dispenses of 50nL (100nL total) 2x NEBNext High-Fidelity 2X PCR Master Mix (NEB, M0541L). The chip was sealed and spun down at 2250xg for 3 mins after each dispense. The PCR on the chip was performed with the following protocol: 5 min at 72 °C and 2 min at 98 °C followed by 15 cycles of 10 s at 98 °C, 30 s at 60 °C, and 5 s at 72 °C, with a final extension at 72 °C for 1 min.

### Quality Control (ICELL8)

The ICELL8 has a built-in imaging system which filters out wells that do not contain a single cell. Thus, empty wells without cells, with more than one single cell, and with doublets, are removed. Subsequently, we filtered single cells with fewer than 100 unique fragments to remove spurious barcodes that can be attributed to an overflow of dispensed PCR material.

A drawback of leveraging a hyperactive transposon in a fusion enzyme to target specific chromatin compartments is that the Tn5 has a high binding affinity for accessible chromatin, the basis of ATAC-seq. Previously, it was shown that this artifact is highly dependent on the concentration of salt in subsequent washes post fusion enzyme binding ^3^. To identify whether our single-cell samples exhibited this artifact, we mapped the percent of reads in each single cell that fell into H3K27me3, H3K4me2, or ATAC specific peaks (Supplementary Fig. 1c). The degree in which repressive H3K27me3 marked chromatin and active accessible chromatin ATAC-seq signal overlapped was minimal as expected whereas an active mark, H3K4me2, had a higher degree of overlap with ATAC-seq data. Correlations of aggregate versus bulk profiles across the 5 kb genome tiles show similar results (Supplementary Fig. 1b).

As an initial test, we wanted to evaluate the robustness of scCUT&Tag by comparing it to scATAC-seq. Therefore, we chose the histone modification K4me2 which was shown to provide similar output to ATAC-seq. A representative genomic track comparing bulk, aggregate, and single cell profiles for K4me2 in H1 and K562 cells, reveal the high-quality resulting data (Supplementary Fig 1a). A low-dimensional embedding, UMAP, clearly separate K562 cells (n=807) from hESC (n=317) (Supplementary Fig. 1d). Projections of published scATAC-seq data (GSE99172) onto our scCUT&Tag embedding align with cell-type specific clusters (Supplementary Fig. 1e).

### Chromatin Profiling: scCUT&Tag using the 10X Genomics system

CUT&Tag was performed with an anti-H3K27me3 antibody (CST#9733) or anti-H3K27ac (MABE647) with 1 million cells as published^3^. Adaptation to the 10X workflow was performed as follows: For all samples except PBMC mixing experiment, the nuclei were spun down at 600g for 3 minutes after the pA-Tn5 binding step. After counting, they were resuspended in 1X Diluted Nuclei Buffer at 2500 nuclei/ul. The nuclei were then prepared for transposition per as per the 10X genomics single cell ATAC-seq protocol (SingleCell_ATAC_ReagentKits_v1.1_UserGuide_RevD). All steps beginning with 1.1 ‘Prepare Transposition Mix’ were performed according to 10X Genomics standard protocol. Libraries were sequenced using an Illumina NovaSeq 6000.

For PBMC mixing experiment, the nuclei were tagmented in high salt (300 mM) as per published protocol^3^. After tagmentation, bovine serum albumin was added to a final concentration of 1%, nuclei were centrifuged at 600 g for 3 mins and then resuspended in 1X Diluted Nuclei Buffer (10X Genomics, PN-2000207) at 2500 nuclei/µL. The 10X genomics single cell ATAC-seq protocol (SingleCell_ATAC_ReagentKits_v1.1_UserGuide_RevD) was used with the following modifications. For step 1.1 ‘Prepare Transposition Mix’, 7 µl ATAC buffer, 3 µl low TE buffer (10 mM Tris pH 8.0, 0.1 mM EDTA) and 5 µl stock nuclei solution were mixed together, omitting the ATAC enzyme as tagmentation had already been performed. All remaining steps beginning with Step 2.0 ‘GEM generation and barcoding’ were performed according to 10X Genomics standard protocol. Libraries were sequenced using an Illumina NovaSeq 6000.

### Data processing

Illumina .bcl files were demultiplexed and converted to fastq format using the cellranger mkfastq function. Resulting fastq files were aligned to the hg38 genome, filtered for duplicates and counted using cellranger atac. An output BED file of filtered fragment data containing the cell barcode was then read into ArchR^7^ as fragment counts in 5kb genome windows which was used in all dimensionality reduction steps across all experiments. We used the ArchR^7^ gene activity score to calculate our CSS as described above. We used LSI dimensionality reduction^7^ using a TFIDF normalization function^25^, UMAP^26^ low dimensional embedding, and clustering using a nearest neighbor graph^25^ performed on data in LSI space.

As the cell line/differentiation experiments used the ICELL8 platform, we did not remove multiplets as this platform uses microscopic imaging to ensure single-cell capture. For droplet partitioning data, we used the following methods to ensure data quality: 1) We first visualized fragment length distribution across clusters. We identified 3 clusters with nucleosomal banding distribution that was consistent with untethered transposition events (Supplementary Fig 3b). 2) We then removed two clusters with high mean fragment counts. 3) We iteratively removed clusters which exhibited non-specific CSS. We accomplished this by calculating CSS significance across clusters using ArchR^7^. Any cluster that did not have any genes that were significantly over-represented or under-represented using significance thresholds of fdr < 0.01 and absolute fold-change > 3 was removed. Bulk projection of down-sampled ChIP-seq data was performed as follows. Raw sequence data aligned to hg38 (BAM files) were downloaded from ENCODE^18^. Data was processed using ChomVAR^27^ by counting reads in 5kb tiled genomes and subsequently used in the bulk projection function in ArchR. Single cell projection was performed using a modified ArchR projection function which did not perform any manipulation of the input data prior to projection. Marker regions/genes for each group were calculated using the getMarkerFeatures function in ArchR. Preranked GSEA (fgsea^28^) was performed using the entire list of marker genes ranked by −log10(pvalue)/sign(foldchange) with the complete MSigDB^29^ set of gene lists. Peak set from scATAC-seq data (see below) was used as a custom annotation set and motif deviations were calculated using addDeviationsMatrix function in ArchR. Pseudotime trajectory was assigned with T1 as a root and Clusters T2 and T4 as an endpoint.

To perform variant calling, we first merged bam output from cellranger-atac using a custom script (https://github.com/scfurl/mergeBams). We then used souporcell^11^ on the merged bam invoking the ‘no_umi’ and ‘skip_remap’ options. Sparse mixture model output from souporcell was log-normalized and colored by the genotype assignment.

### Quality control and data processing for brain tumor ATAC-seq

Nuclei preparation from snap frozen brain tumor tissue was performed as described above and standard single cell ATAC-seq workflow was performed as per manufacturer guidelines (10X Genomics). Sequencing data was processed using the cell ranger atac package. An output BED file of filtered fragment data containing the cell barcode was then read into ArchR^7^ using 500 bp genome windows. We used LSI dimensionality reduction^7^ using a TFIDF normalization function^25^, UMAP^26^ low dimensional embedding, and clustering using a nearest neighbor graph^25^ performed on data in LSI space. Tumor cells were identified as the largest cluster containing high gene activity scores for marker genes SOX2 and PTPRZ1. This cluster was used for peak calling using the MACS2 wrapper in ArchR with standard parameters.

### External Data

Data from the following identifiers were downloaded from the ENCODE portal (https://www.encodeproject.org“) and Gene Expression Omnibus (https://www.ncbi.nlm.nih.gov/geo/). For figure 1b and supplementary figure 1a, 1f, and 1g: GSE124557. For figure 1d and supplementary figure 2c: GSE75748. In addition, for the purposes of this study hESC differentiated timepoint 1.5 (scRNA-seq) is approximated to be day 2 in the GSE75748 dataset. For supplementary figure 2b, 2c, and 2e: GSE99172, GSE99173, GSE124557, and GSE85330. For figure 2b: ENCSR000ASK, ENCSR043SBG, ENCSR103GGR, ENCSR404MOX, and ENCSR939JZW. For figure 3c, the following data sets were used: ENCFF363TCY, ENCFF911MNN.

## Data Availability

Sequencing data are deposited in the Gene Expression Omnibus (GEO) with accession code (pending). There are no restrictions on data use.

## Code Availability

Code used in this study can be found on Github at https://github.com/Henikoff/scCUT-Tag.

## Acknowledgements

We thank Eric Holland and members of the Holland lab for providing shared space for experimental work. We also thank Michael Meers, Derek Janssens, Manu Setty, Jitendra Thakur, and other members of the Henikoff lab for helpful suggestions, discussions, and mentorship. We thank BioRender helping us create figures. This work was supported by the Howard Hughes Medical Institute (S.H.), grants R01 HG010492 (S.H.) and R01 GM108699 (K.A.) and K08 CA245037 (P.J.C.) from the National Institutes of Health, an HCA Seed Network grant from the Chan-Zuckerberg Initiative (S.H., A.P.P, S.N.F, R.G., K.A., Y.Z.), a Burroughs Wellcome Career Award for Medical Scientists (A.P.P) and an American Cancer Society Mentored Scholar Award (S.N.F).

## Author Contributions

S.J.W, S.N.F, A.B.M, H.K-O, A.H.F, S.N.E, and J.F.S processed samples and performed experiments. S.J.W, S.N.F, Y.Z., R.G and A.P.P performed and/or provided input on data processing and analysis. K.C, P.J.C, and C.D.K provided access to tissue samples and assisted with processing. S.J.W, S.N.F, K.A., S.H, and A.P.P wrote the manuscript with input from all authors.

## Competing Interests

S.N.F has received research support from Lyell Immunopharma. R.G. has received consulting income from Juno Therapeutics, Takeda, Infotech Soft, Celgene, Merck and has received research support from Janssen Pharmaceuticals and Juno Therapeutics, and declares ownership in CellSpace Biosciences. H.S.K and S.H have filed patent applications related to this work.

## Notes

### Competing Interest Statement

The authors have declared no competing interest.

### Summary of Updates

We have prepared a revised manuscript which incorporated biologic and technical replicates of H3K27me3 data and from a second histone mark (K27ac) in PBMCs. We have also provided deeper insight into data quality of the tumor samples.

