## Supplementary Material for "Single-cell analysis of chromatin silencing programs in development and tumor progression"

a PBMCs

| Cluster (Cell Type) | H3K27me3 | H3K27ac | Normal adult ranges |
|---|---|---|---|
| **Mono** | **20.90%** | **47.60%** | **10-30%** |
| **NK or T** | **66.40%** | **48.30%** | **45-70% T**<br>**5-10% NK** |
| **B** | **12.70%** | **4%** | **5-15%** |

b **Glioblastoma**

| | scATAC-seq | scCUT&Tag |
|---|---|---|
| **Microglia** | **20.70%** | **21.50%** |
| **Astrocytes** | **2.70%** | **3.50%** |
| **Tumor** | **50.40%** | **57.50%** |
| **Oligodendrocytes** | **4.40%** | **5.40%** |
| **Neurons** | **19.00%** | **12.10%** |

Supplementary Table 1: a) A table comparing the proportions of major cell types in PBMCs identifed by scCUT&Tag for H3K27me3 and H3K27ac to scATAC-seq and FACS. b) A table comparing the proportions of major cell types identified in the glioblastoma sample by scCUT&Tag versus scATAC-seq.

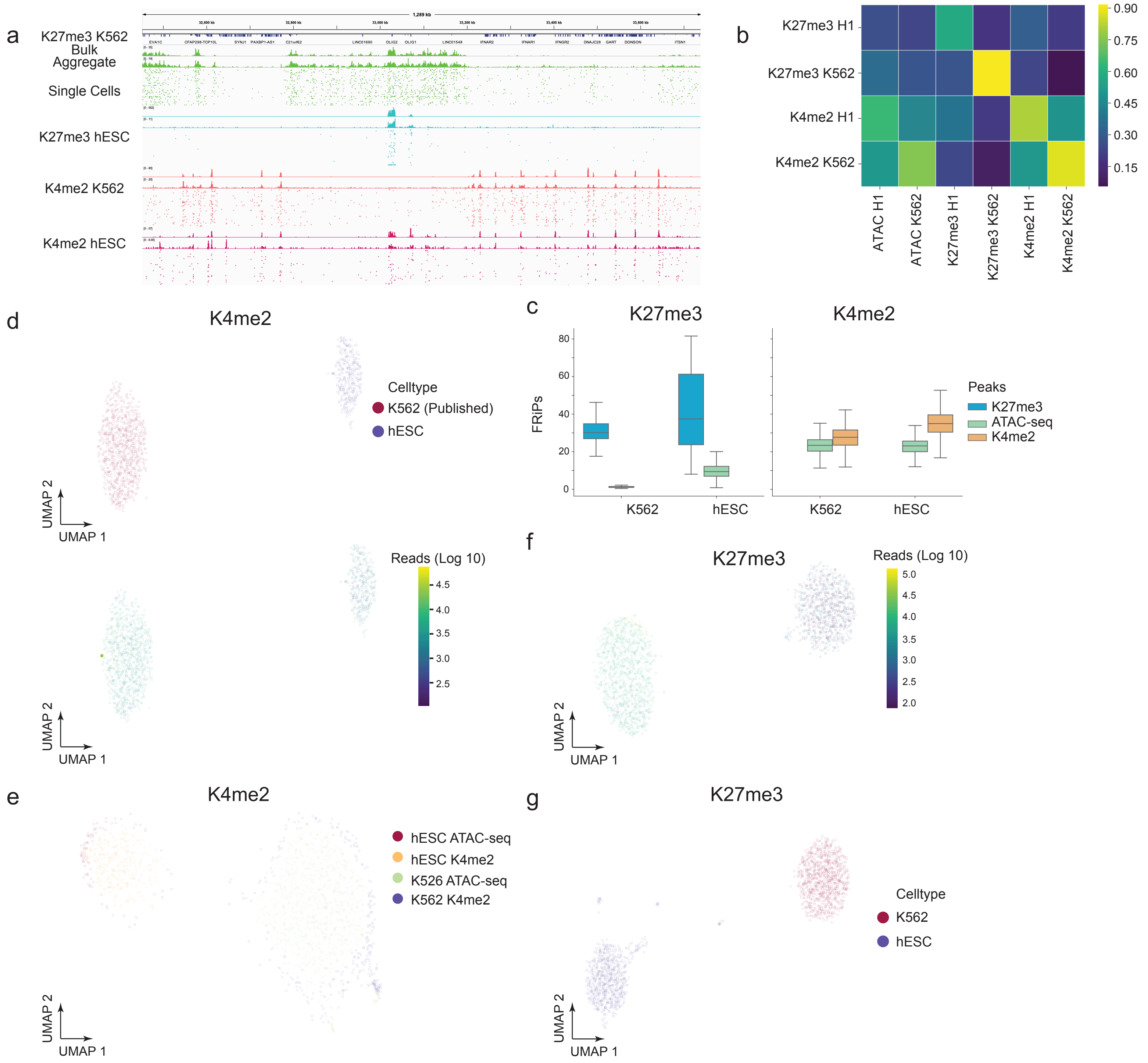


Supplementary Figure 1: A) Representative chromatin landscapes across 1.2 Mb segment of the human genome for bulk, aggregate, and the top 50 single cells. B) A heatmap comparing the Pearson correlation of fragment counts in 5 kb bins across the genome for bulk and aggregate datasets. C) Boxplots illustrating the fraction of reads in peaks (FRiPs) for each single cell from peaks called in bulk datasets. ATAC-seq datasets are used as a control for background. D) UMAP embedding of single cells for H3K4me2 shaded by their respective celltype (top) and unique number of reads bottom). E) UMAP embedding of singles cells for H3K4me2. Published single cell ATAC-seq data from the same cell type are taken and projected onto the H3K4me2 low dimensional space. F) UMAP projection of H3K27me3 single cells shaded with the number of unique fragments per cell. G) UMAP projection of K27me3 single cells for K562 and hESC when K562 are downsampled to 365 reads per cell.

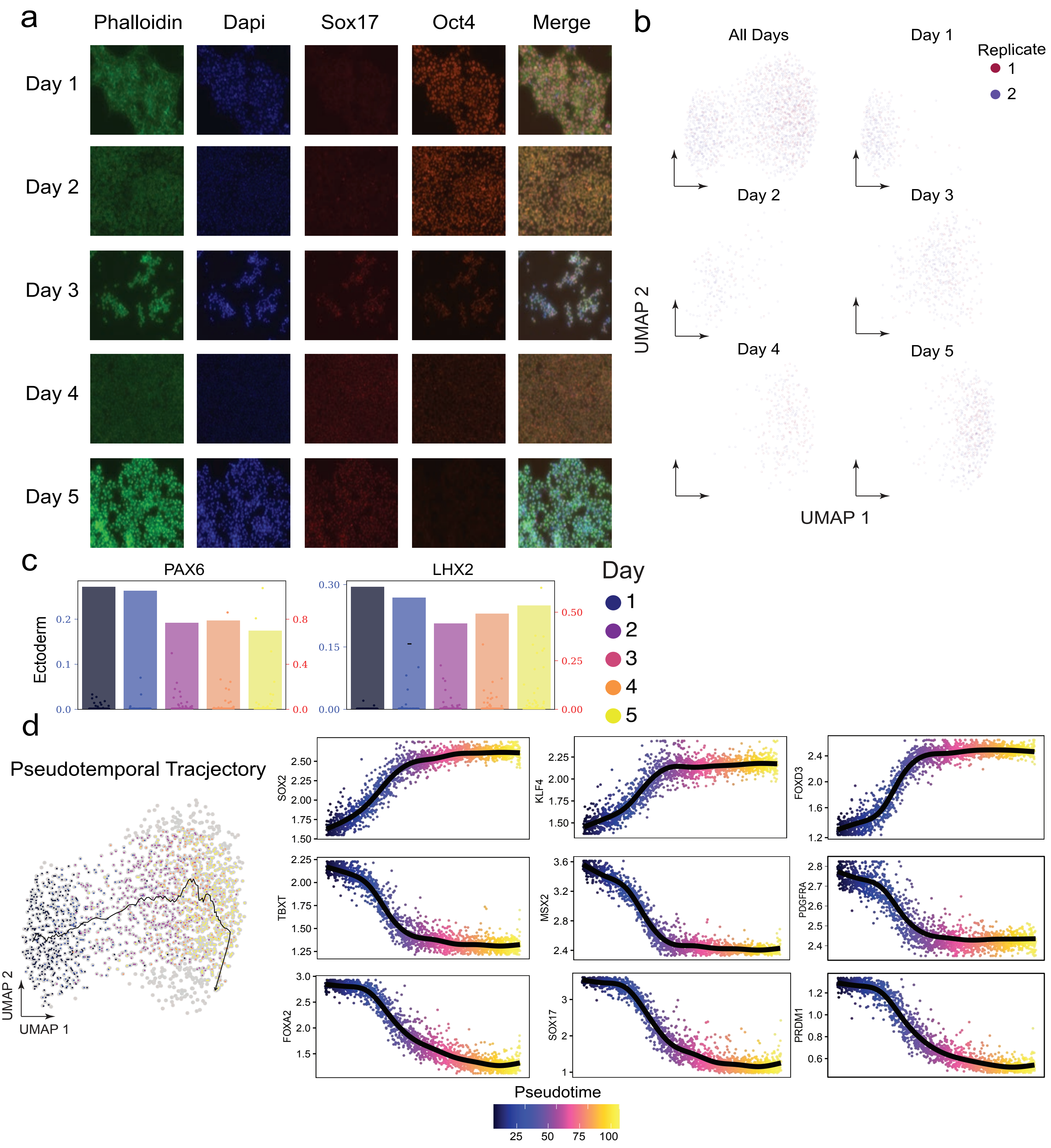


Supplementary Figure 2: a) Immunofluorescence, conducted as shown in for well known TF markers that indicate either pluripotency (Oct4) or definitive endoderm (Sox17). b) A grid plot illustrating UMAP projections for two technical replicates along different time points. c) A bar plot representing the percent of single cells that are repressed at each specific gene. The superimposed jitters depict scRNA-seq for the same timepoint. The left axis corresponds to scCUT&Tag (percent of single cells repressed) and right corresponds to scRNA-seq (normalized mRNA counts).d) On the right a UMAP embedding showing the pseudotemporal ordering of single cells from a hESC to endoderm time course. On the left, dot plots of pseduo-time versus imputed chromatin silencing score at the gene of interest.

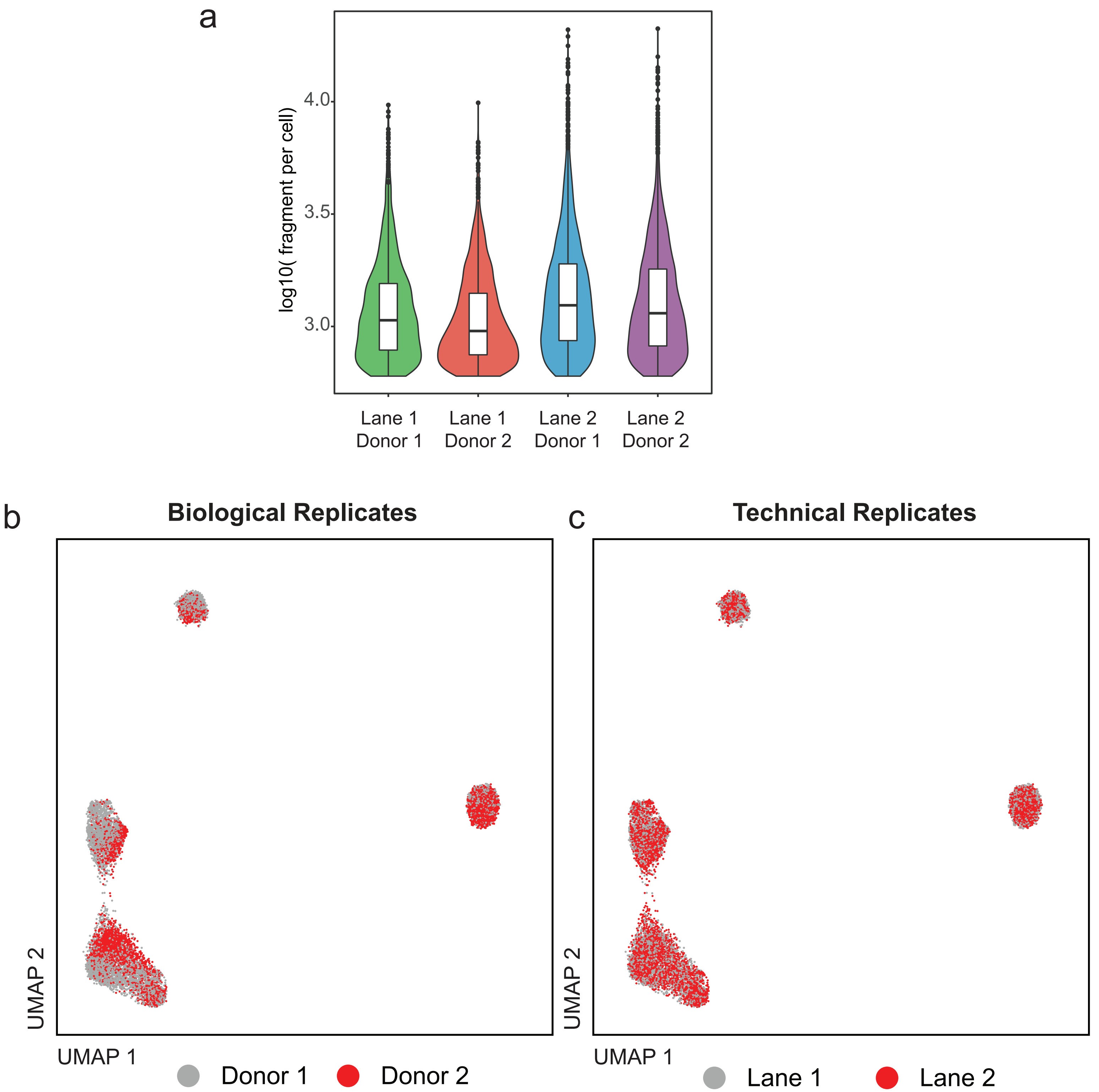


Supplementary Figure 3 - scCutTag biologic replicates (across donors) and technical replicates (across lanes).
A. Violin plots depicting number of unique fragments in each cell within different biologic and technical replicates. Number of fragments recovered is similar across either different donors or across different 10X lanes.
B. UMAP embedding colored by biologic replicates. Donors 1 and 2 overlap demonstrating minimal donor specific effects on clustering.
C. UMAP embedding colored by technical replicates. Pooled PBMCs were run on two separate lanes of a 10X microfluidic chip. The two technical replicates overlap without replicate specific effects on clustering.

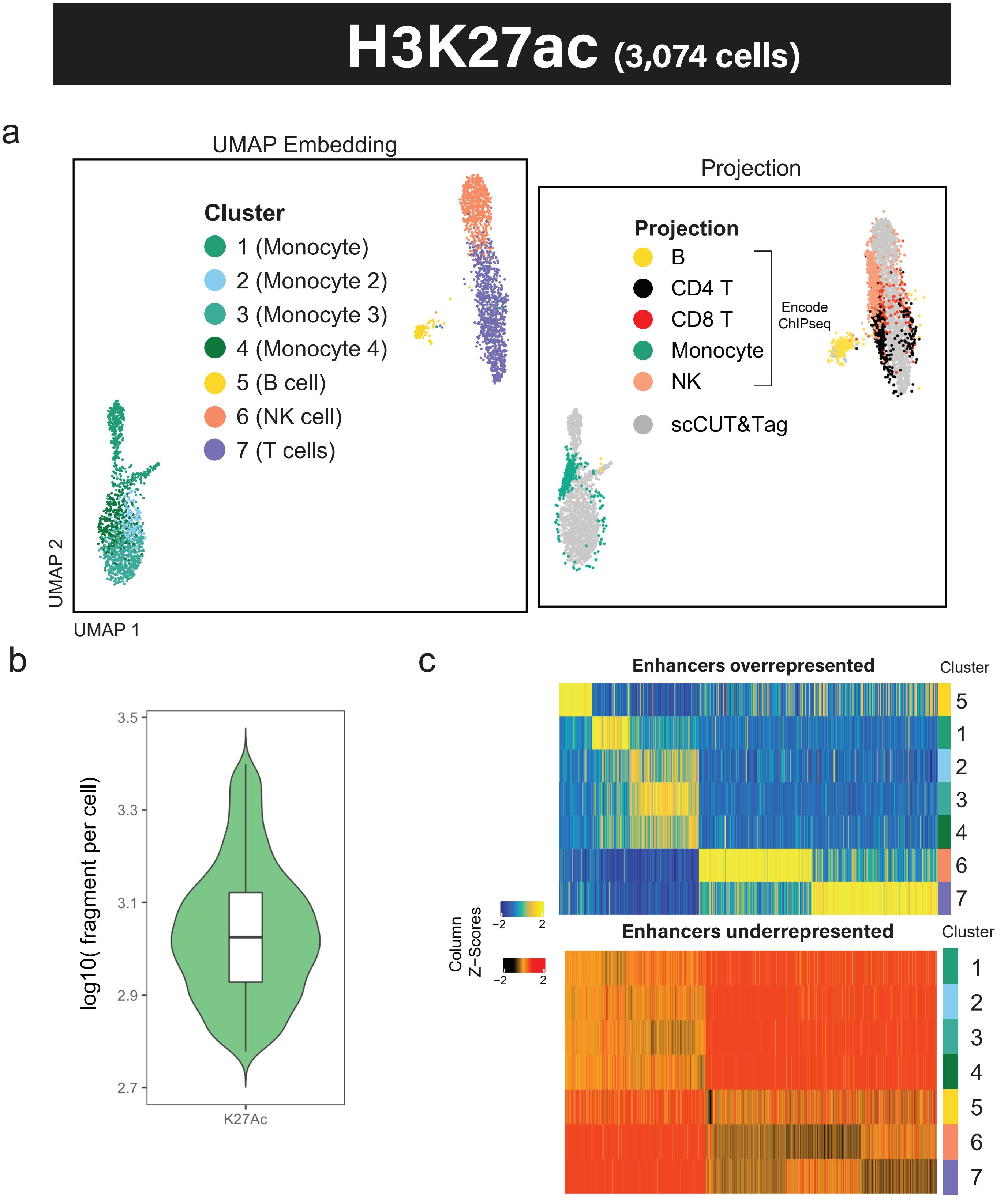


Supplementary Figure 4: scCUT&Tag for H3K27ac readily identify major subtypes in PBMC
a) Left - UMAP embedding of single cell data from PBMC. Unsupervised clustering revealed 7 clusters. Right - UMAP projection of downsampled ChIPseq bulk data from primary sorted bulk datasets for major PBMC cell-types on single cell CUT&Tag data on left. ENCODE accessions: ENCSR000ASJ - Monocytes, ENCSR000AUP - B cells, ENCSR007HLH - CD8 T, ENCSR138DOM - CD4, ENCSR391EQV - NK. b) Violin plot of fragments per cell. c) Heatmap of genes with significantly high (top) or low (bottom) H3K27ac signal in each cluster (row). Fold change > 2 (top) or < -2 (bottom); q-value < 0.05 (both).

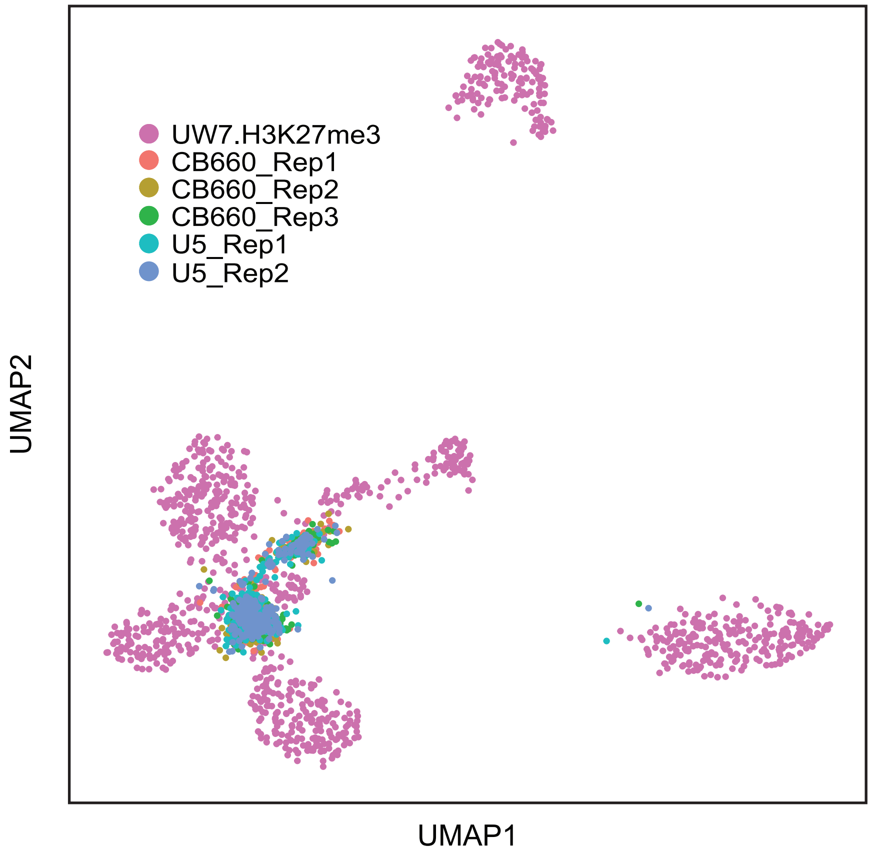


Supplementary Figure 5: Projection of published bulk Cut&Run data for two neural stem cell lines CB660 (three replicates) and U5 (two replicates) onto the single cell patient data. Neural stem cells localize to the center of the tumor cell cluster but also adjacent to the astrocyte cluster.

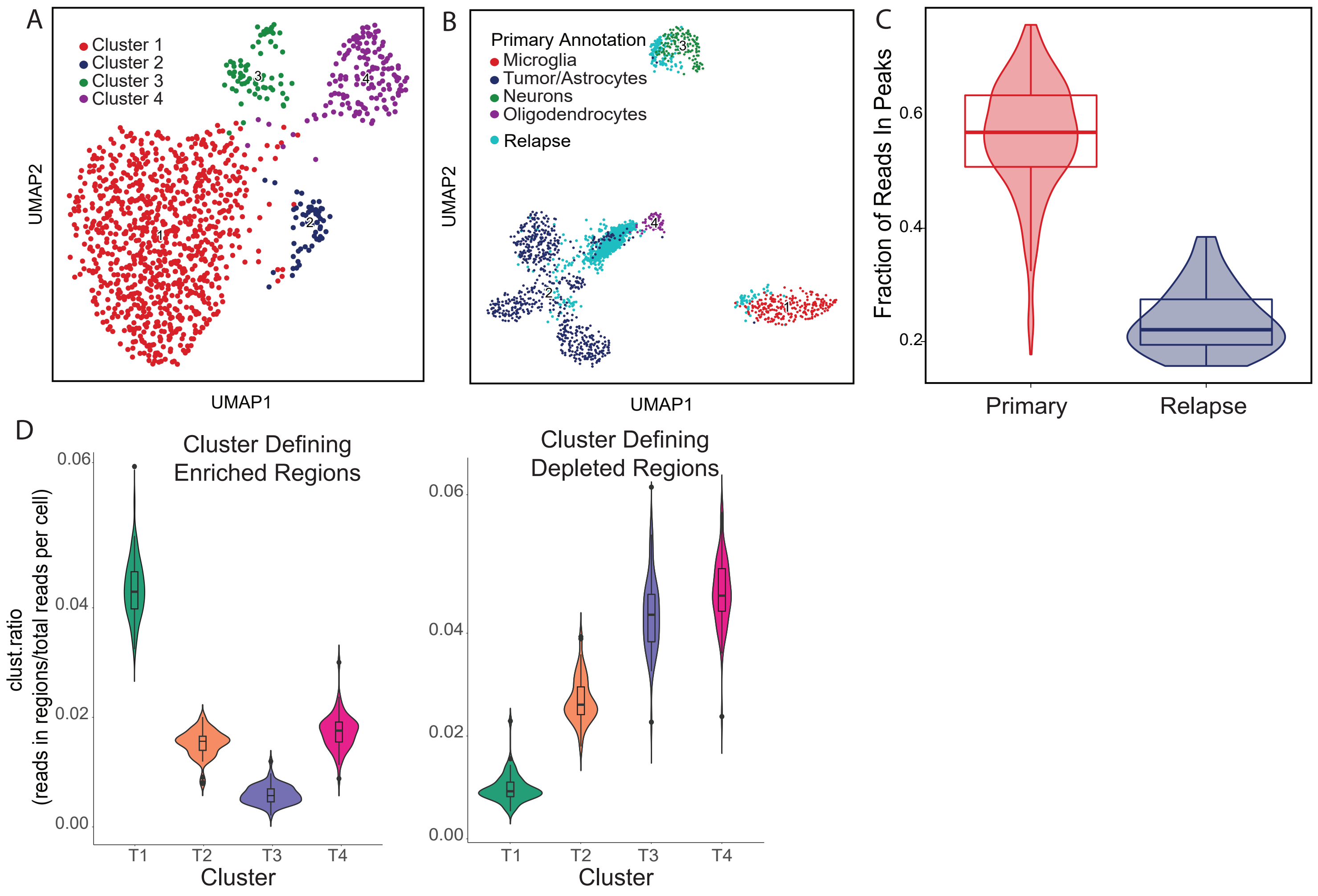


Supplementary Figure 6: Identification of tumor cells from relapsed sample.
A: UMAP embedding of relapse sample from patient (n=1168 cells) based on H3K27me3 signal. Four clusters are identified.

B: Projection of single cells from relapsed sample onto primary patient sample using UMAP transform allows identification of the four clusters in the relapse data corresponding to microglia, tumor cells, astrocyes, and neurons. The majority of the cells colocalize with the astrocyte cluster from the primary tumor. 71 cells colocalized with the tumor cell cluster. There are several reasons why recovery of tumor cells was relatively low. Due to the nature of the sampling done at autopsy we could not ensure that we had sampled regions with high tumor cell density. In addition, post treatment tumors are often characterized by increases in stromal cells, particularly reactive astrocytes.

C: Increased background in relapse sample. We called peaks using HOMER with settings appropriate for broad domains (see methods) on the aggregate sequencing data for each sample and then calculated the fraction of reads in peaks (FRiP) on a per cell basis. The value for each cell is plotted. Overall, the primary sample had a higher (mean 0.57 versus 0.22) FRiP value. This suggested a higher amount of background signal in the post mortem specimen.

D: Violin plots demonstrating similarity of autopsy cells to cluster 1 from the primary tumor. Genomic regions that most significantly (FDR<0.05, log2FC>1 or log2FC<1) distinguished tumor clusters based on chromatin silencing score were calculated. Cluster ratio was calculated by counting reads in each autopsy cell in cluster defining regions and dividing by total number of unique fragments for that cell. Distributions were plotted for each set of cluster defining regions (T1-T4). This was perfromed for regions that define each cluster by both enrichment (positive, left panel) and depletion (negative, right panel). Violin plots demonstrate that autopsy cells are most similar to T1 based on both enrichment and depletion of reads in T1 definining regions of the genome.

A

Postively Enriched Gene Sets

| Pathway | Gene ranks | NES | pval | padj |
|---|---|---|---|---|
| GO_OLFACTORY_RECEPTOR_ACTIVITY | | 2.47 | 1.5e−31 | 2.8e−27 |
| REACTOME_OLFACTORY_SIGNALING_PATHWAY | | 2.44 | 1.5e−30 | 1.4e−26 |
| GO_SENSORY_PERCEPTION_OF_SMELL | | 2.40 | 5.6e−30 | 3.5e−26 |
| GO_SENSORY_PERCEPTION_OF_CHEMICAL_STIMULUS | | 2.24 | 4.3e−24 | 1.3e−20 |
| GO_DETECTION_OF_STIMULUS_INVOLVED_IN_SENSORY_PERCEPTION | | 2.19 | 4.2e−22 | 1.1e−18 |
| KEGG_OLFACTORY_TRANSDUCTION | | 2.24 | 9.3e−21 | 1.9e−17 |
| REACTOME_G_ALPHA_S_SIGNALLING_EVENTS | | 1.98 | 1.8e−16 | 2.9e−13 |
| KEGG_ASCORBATE_AND_ALDARATE_METABOLISM | | 2.11 | 1.4e−09 | 9.4e−07 |
| REACTOME_GLUCURONIDATION | | 2.06 | 5.2e−08 | 2.5e−05 |
| KEGG_PENTOSE_AND_GLUCURONATE_INTERCONVERSIONS | | 2.05 | 7.0e−08 | 3.1e−05 |
| GO_ODORANT_BINDING | | 2.06 | 3.7e−07 | 1.3e−04 |
| GO_CELLULAR_GLUCURONIDATION | | 1.99 | 7.4e−07 | 2.3e−04 |
| GO_GLUCURONOSYLTRANSFERASE_ACTIVITY | | 2.03 | 1.1e−06 | 3.3e−04 |
| KEGG_PORPHYRIN_AND_CHLOROPHYLL_METABOLISM | | 2.05 | 1.4e−06 | 3.6e−04 |
| GO_URONIC_ACID_METABOLIC_PROCESS | | 1.97 | 1.6e−06 | 4.0e−04 |
| MODULE_505 | | 1.88 | 1.5e−05 | 2.3e−03 |
| VERHAAK_GLIOBLASTOMA_PRONEURAL | | 1.70 | 3.9e−05 | 4.9e−03 |
| GO_UDP_GLYCOSYLTRANSFERASE_ACTIVITY | | 1.69 | 1.6e−04 | 1.3e−02 |
| MORF_MAP3K14 | | 1.73 | 1.9e−04 | 1.4e−02 |
| KEGG_STEROID_HORMONE_BIOSYNTHESIS | | 1.82 | 3.0e−04 | 2.0e−02 |

B

Negatively Enriched Gene Sets

| Pathway | Gene ranks | NES | pval | padj |
|---|---|---|---|---|
| GO_SENSORY_ORGAN_DEVELOPMENT | | −1.80 | 3.4e−10 | 2.4e−07 |
| GO_CENTRAL_NERVOUS_SYSTEM_NEURON_DIFFERENTIATION | | −2.05 | 2.7e−10 | 1.9e−07 |
| GO_FOREBRAIN_DEVELOPMENT | | −1.88 | 1.4e−10 | 1.0e−07 |
| GO_ANTERIOR_POSTERIOR_PATTERN_SPECIFICATION | | −2.04 | 1.3e−10 | 9.9e−08 |
| MIKKELSEN_MEF_HCP_WITH_H3K27ME3 | | −1.78 | 7.5e−11 | 6.1e−08 |
| GO_EMBRYONIC_ORGAN_DEVELOPMENT | | −1.91 | 7.0e−12 | 5.9e−09 |
| GO_GLAND_DEVELOPMENT | | −1.91 | 4.7e−12 | 4.2e−09 |
| GO_SKELETAL_SYSTEM_DEVELOPMENT | | −1.87 | 1.6e−12 | 1.5e−09 |
| MEISSNER_NPC_HCP_WITH_H3K4ME2_AND_H3K27ME3 | | −1.99 | 1.6e−12 | 1.5e−09 |
| GO_EMBRYONIC_ORGAN_MORPHOGENESIS | | −2.05 | 8.0e−13 | 8.4e−10 |
| MIKKELSEN_NPC_HCP_WITH_H3K27ME3 | | −2.00 | 2.8e−13 | 3.1e−10 |
| NIKOLSKY_BREAST_CANCER_17Q21_Q25_AMPLICON | | −2.03 | 2.4e−13 | 2.8e−10 |
| GO_CELL_FATE_COMMITMENT | | −2.11 | 7.1e−14 | 8.8e−11 |
| MIKKELSEN_MCV6_HCP_WITH_H3K27ME3 | | −1.97 | 6.9e−14 | 8.8e−11 |
| GO_DNA_BINDING_TRANSCRIPTION_ACTIVATOR_ACTIVITY | | −2.03 | 4.9e−16 | 7.0e−13 |
| GO_EMBRYONIC_MORPHOGENESIS | | −2.02 | 5.0e−19 | 8.5e−16 |
| GO_CIS_REGULATORY_REGION_BINDING | | −2.02 | 3.1e−19 | 5.8e−16 |
| GO_REGIONALIZATION | | −2.21 | 4.4e−21 | 1.0e−17 |
| GO_PATTERN_SPECIFICATION_PROCESS | | −2.23 | 3.5e−25 | 1.3e−21 |
| MEISSNER_BRAIN_HCP_WITH_H3K27ME3 | | −2.41 | 7.5e−28 | 3.5e−24 |
| | 0 5000 10000 15000 20000 | | | |

Supplementary Figure 7: Gene set enrichment analysis of cluster 1, which is highly enriched in the relapse sample. Gene set enrichement was performed by performing a pairwise test for CSS at each gene in cluster 1 versus all other clusters. Differentially silenced genes were then rank ordered using -log10(pval)/sign(log2FC) as a metric and all MSigDB gene sets were interrogated.

A: Positively enriched gene sets: genesets in this group demonstrate high PcG activity at their gene loci, and therefore represent silenced programs. The Verhaak_glioblastoma_proneural geneset is positively enriched for PcG mediated silencing.

B: Negatively enriched gene sets: genesets inthe group demonstrated low PcG activity at their gene loci. The most negatively enriched term is whole brain H3K27me3 profiles from Meissner et.al. Given that whole brain represents most differentiated cell types (neurons), it is consistent that genes in this geneset would be negatively enriched for PcG given that glioblastoma cells are not terminally differentiated.rzsa11A11a

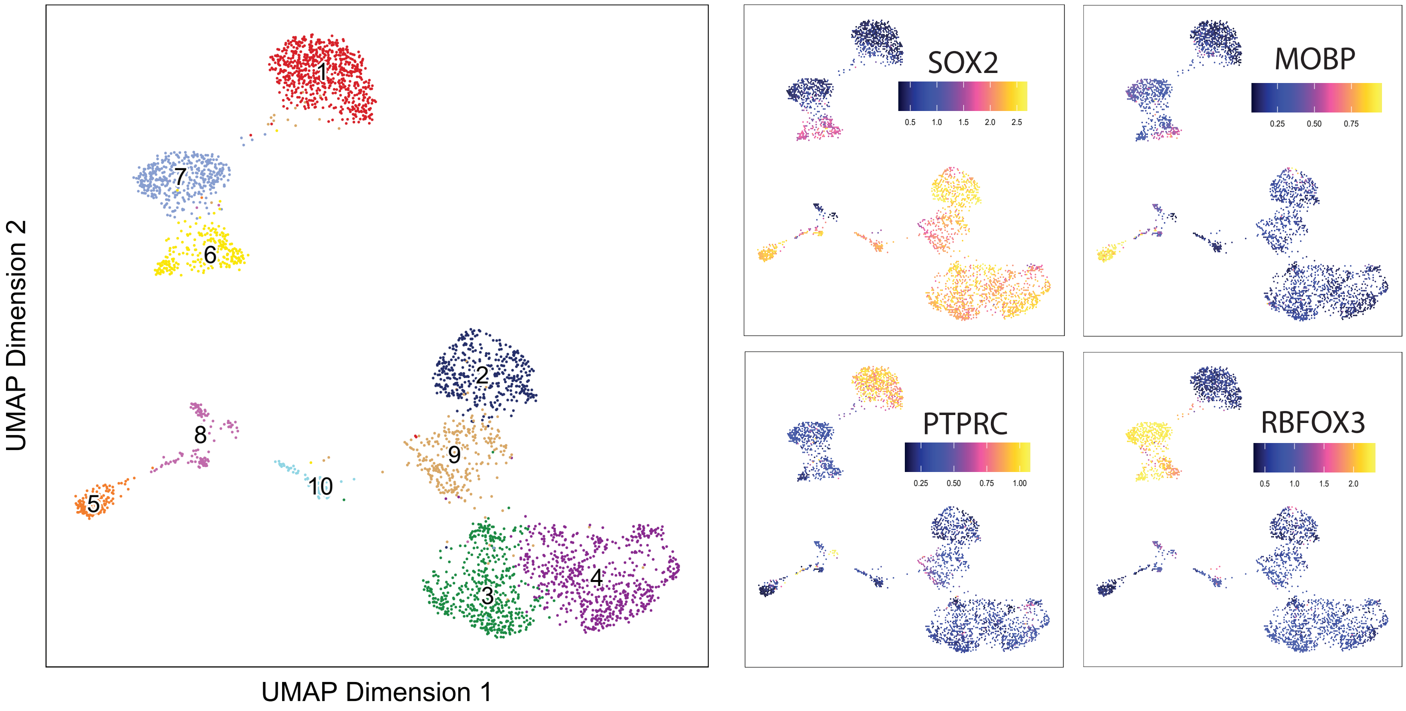


Supplementary Figure 8: Identification of accessible chromatin peaks in scATAC-seq data.

A. UMAP embedding of single cell ATAC-seq data from patient UW7. 3541 single cells were profiled and dimensionality reduction and clustering were performed (see methods). 10 individual clusters are identified.

B. UMAP plots colored by gene activity scores for individual marker genes for each large cluster that were identified based on gene scores that were most different across the clusters. SOX2 identifies cells of neural lineage, MOBP identifies oligodendrocytes, PTPRC identifies microglia, and RBFOX3 identifies neurons. The largest subcluster of the SOX2 positive group was identified as tumor cells based on top differentially expressed genes based on gene activity scores (PTPRZ1, PDGFRA, EGFR). Cells in the tumor cell clusters were aggregated and peaks were called using MACS2, identifying accessible enhancers and promoters which were used for analysis of motif silencing in the H3K27me3 dataset.
